# Real world data based evaluation of a novel target-mediated drug disposition approximation model

**DOI:** 10.64898/2026.01.20.700502

**Authors:** Hyeseon Jeon, Woojin Jung, Hwi-yeol Yun, Soyoung Lee, Jae Kyoung Kim, Jung-woo Chae, Jong Hyuk Byun

**Author notes:** Corresponding author E-mail: hy.yun, s.lee, jk.kim, jw.chae jh.byun. These authors contributed equally to this work as first authors (Hyeseon Jeon, Woojin Jung). These authors contributed equally to this work as corresponding authors.

## Abstract

Target-mediated drug disposition (TMDD) models have been widely used to describe nonlinear pharmacokinetic profiles driven by high-affinity, low-capacity drug–target binding. A pTMDD model, derived by applying the Padé approximation of the quasi-steady-state (QSS) model (qTMDD) was previously proposed. Although pTMDD model showed a comparable estimation accuracy while maintaining computational efficiency, further validation in realistic clinical scenarios and comprehensive performance evaluations have been needed to assess its practical applicability. Here, we evaluated the pTMDD model using five clinical datasets and extended the previous study that focused on simulations. Using the full TMDD as a reference, the approximation models were compared in terms of the parameter estimation results (parameter estimates, relative standard error values and model diagnostics) and computational efficiency (estimation and bootstrap runtimes). The pTMDD model, previously validated in simulation settings, also preserved the estimation accuracy while reducing the computation time of the clinical data. Both pTMDD and qTMDD remained close to the full TMDD model, whereas Michaelis-Menten TMDD (mTMDD) model showed substantial discrepancies especially at low doses, including biased estimates for key TMDD-related parameters (e.g., k_deg_, k_int_, k_rec_, and k_up_) and higher objective function values. Moreover, pTMDD was faster than qTMDD in four of the five cases compared to the full TMDD. The time savings were particularly pronounced for larger datasets, supporting the computational efficiency of pTMDD. Q2PCONV, an R Shiny application that converts NONMEM code from qTMDD to pTMDD, was also developed, thereby making this new approximation more accessible to researchers. The findings support pTMDD as a practical alternative to existing TMDD approximation models.

**Author Summary:** Target-mediated drug disposition (TMDD) models describe a high-affinity, low-capacity binding between drug and its target. To avoid overparameterization, approximation models have been used. The two primary models are Michaelis-Menten model (mTMDD), which is accurate only at high doses, and Quasi-steady-state (qTMDD), which is accurate in wider ranges but requires longer runtime. We have proposed a new approximation model named pTMDD. Here, we evaluated pTMDD using five real clinical trial datasets to assess its practical usefulness. pTMDD produced parameter estimates closer to those from the full TMDD model and showed lower uncertainty than both the full TMDD and mTMDD models. In terms of computational efficiency, pTMDD reduced estimation time by an average of 11% and bootstrap time by an average of 6% relative to qTMDD across cases. In addition, we also developed an R shiny application to help researchers apply pTMDD in practice. Our work supports pTMDD as a practical and efficient tool for TMDD modeling in drug development.

## Introduction

Target-mediated drug disposition (TMDD) is a nonlinear pharmacokinetic phenomenon caused by the high-affinity binding of a compound to its pharmacological targets [1]. TMDD models explicitly describe drug–receptor binding and dissociation, degradation of unbound drugs, and internalization and degradation of the drug–receptor complex. Thus, they account for dose-dependent behavior [2]. At high doses, the drug rapidly binds to and saturates the receptor, and most of the drug is then eliminated through traditional linear pathways. However, at low doses, the receptor is not saturated and a large fraction of the drug is cleared via receptor-mediated endocytosis of the drug–receptor complex [3]. By capturing nonlinear pharmacokinetics that cannot be adequately described by compartmental models, TMDD models have become increasingly important, particularly with the growing development of biological pharmaceutics.

TMDD models involve many parameters and are highly complex, often leading to instability when data availability is limited [4]. Various approximation methods have been developed to address this problem. The quasi-equilibrium (QE) model by Mager and Krzyzanski (2005) [5] was the first TMDD approximation, followed by the quasi-steady-state (QSS) and Michaelis– Menten (MM) models by Gibiansky et al. (2008) [6]. Of these, the QSS model is the most widely used; however, its equations are complex and often require long computation times for parameter estimation. The MM model is simpler and more computationally efficient. However, its accuracy can be poor under low doses.

To address these limitations, a new approximation model was previously derived by applying the Padé approximation to the QSS model (named pTMDD). The pTMDD model was valid regardless of initial drug concentration(C_0_) whereas mTMDD was valid only at high C_0_, indicating its estimation accuracy. The pTMDD model required shorter estimation time compared to qTMDD, indicating its computational efficiency [7]. However, the work was primarily based on simulation studies and was evaluated using only a limited number of clinical datasets.

In the current study, we aimed to extend previous work by evaluating the pTMDD model using five clinical trial datasets, thereby building on earlier evaluations conducted in a simplified setting. This multi-dataset evaluation provides a stronger empirical basis by examining whether pTMDD performs consistently across diverse pharmacological settings. We assessed pTMDD in terms of estimation results and computational efficiency. Instead of comparing only point estimates, parameter estimate results were assessed by comparing parameter estimates and uncertainty using both variance-covariance matrix-based and bootstrap-based summaries. We also broadened the evaluation of computational efficiency by measuring both estimation and bootstrap runtime. In addition, Q2PCONV, an R Shiny application that enables a convenient transition from the qTMDD to the pTMDD model, was developed. By demonstrating that pTMDD provides accurate and efficient parameter estimation in clinical data and by offering user-friendly software, we expect this work to facilitate the broader and more practical use of TMDD modeling in drug development.

## Results

### Classification of five clinical datasets based on validity conditions

Five clinical datasets were selected from clinical trials in which high-affinity binding between the drug and its receptor(target) was observed. These consisted of population TMDD pharmacokinetic (PK) models for the anti–PD-L1 monoclonal antibody [8], anakinra [9], anti-CD47 monoclonal antibody [10], IL-1Ra Fc fusion protein [9], and IL-7 Fc fusion protein [11]. The models were fitted to 396, 93, 211, 472, and 410 time versus plasma concentrations.

In our previous work, we have proposed the simplified validity criterion as follows:

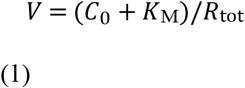

mTMDD provided accurate results only when this value was greater than 1, whereas both qTMDD and pTMDD provided accurate results regardless of this value. To account for the underlying biological process [12], TMDD mediated by FcRn was considered in addition to TMDD involving the pharmacological receptor. Accordingly, *V*_*1*_ and *V*_*2*_were defined:

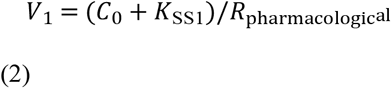

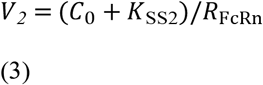

and V values were calculated for each dataset (Table 1).

**Table 1.** Classification of each case according to the V-value.

| **Drug:** | **Case 1)**<br>**anti-PD-L1**<br>**mAb** | **Case 2)**<br>**Anakinra** | **Case 3)**<br>**anti-CD47**<br>**mAb** | **Case 4)**<br>**hIL-1Ra-hyFc** | **Case 5)**<br>**rhIL-7-hyFc** |
| --- | --- | --- | --- | --- | --- |
| $V_1$ : | 611.89 | 23.86 | 152.81 | 21.92 | 21.9 |
| $V_2$ : | 252.37 | - | 7.77 | 5.46 | 0.0003 |
| $C_0$ | 2.14 μM<br>(1.53 μM) | 36 nM<br>(6.68 nM) | 1.91 μM<br>(1.4 μM) | 34.39 nM<br>(29.14 nM)<br>*\*For $V_2$, 3839 nmol (2906 nmol)* | 0.008 nM<br>(0.011 nM) |
| $K_{SS1}$ (receptor) | 0.0016 μM | 0.75 nM | 0.046 μM | 14.5 nM | 29.78 nM |
| $K_{SS2}$ (FcRn) | 71.3 μM | - | 10.6 μMol | 237 nmol | 29.82 nM |
| $R_{\text{pharmacological}}$ | 0.0035 μM | 1.54 nM | 0.0128 μM | 2.23 nM | 1.36 nM |
| $R_{\text{FcRn}}$ | 0.291 μM | - | 26.5 μMol | 746 nmol | 91000 nM |

$C_0$ : Initial drug concentration; $R_{\text{pharmacologic}}$ : total pharmacological receptor concentration; $R_{\text{FcRn}}$ : Total FcRn concentration or amount. For cases 2, 4, and 5 in which the drug was administered subcutaneously or intramuscularly, the mean concentration was used. For Case 4, because the original model approximated drug-FcRn binding based on quantitative equilibrium rather than concentration, with FcRn and KSS2 measured in nmol, the drug dose was used instead of $C_0$ as an approximation.

Based on these indices, the approximation performance was expected to differ by case. In Case 1, where V_1_ ≫ 1, and in Case 2, where the TMDD with FcRn was not required, all approximation models were expected to show results similar to those of the full TMDD model. In contrast, in Case 5, where V_2_ ≪ 1, the mTMDD was anticipated to differ markedly from the full TMDD model, whereas the qTMDD and pTMDD were expected to provide similar results.

### pTMDD outperformed mTMDD in the estimation results

In the parameter estimation step, parameter estimate accuracy, model diagnostics and parameter uncertainty were evaluated. Relative accuracy was assessed based on how closely the parameter estimates matched those from the full TMDD model, and the parameter uncertainty was measured by the RSE. Two approaches were applied in accordance with the methodology described by Jonsson and Nyberg [13]: (1) a parametric method using the variance–covariance matrix, and (2) a nonparametric method using bootstrap results.

Parameter estimate accuracy was better with both pTMDD and qTMDD than with mTMDD under both approaches. In Cases 1–3, where the validity criteria were satisfied for all the approximation models, no substantial differences in the parameter estimates were observed. In Case 4, although V_1_ and V_2_ were both greater than 1, their values were smaller than in the other cases, and mTMDD showed clear estimation failures. Specifically, mTMDD failed to estimate parameters such as *k*_*deg*_, *k*_*int*_, *k*_*rec*_ (parametric approach, Fig 1a), and *Q* (Nonparametric approach, Fig 1b), whereas pTMDD and qTMDD did not show such failures. In Case 5, where V_1_ > 1 but V_2_ < 1, mTMDD failed to estimate additional parameters such as *Q* (parametric approach, Fig 1a), *k*_*rec*_ and *V*_*D*_ (Nonparametric approach, Fig 1b). These failures were not observed with pTMDD or qTMDD. Consistent patterns were also observed in model diagnostics such as OFV(S1 Table), basic goodness-of-fit plots(S1 Fig) and visual predictive check plots(S2 Fig).

**Fig 1.**
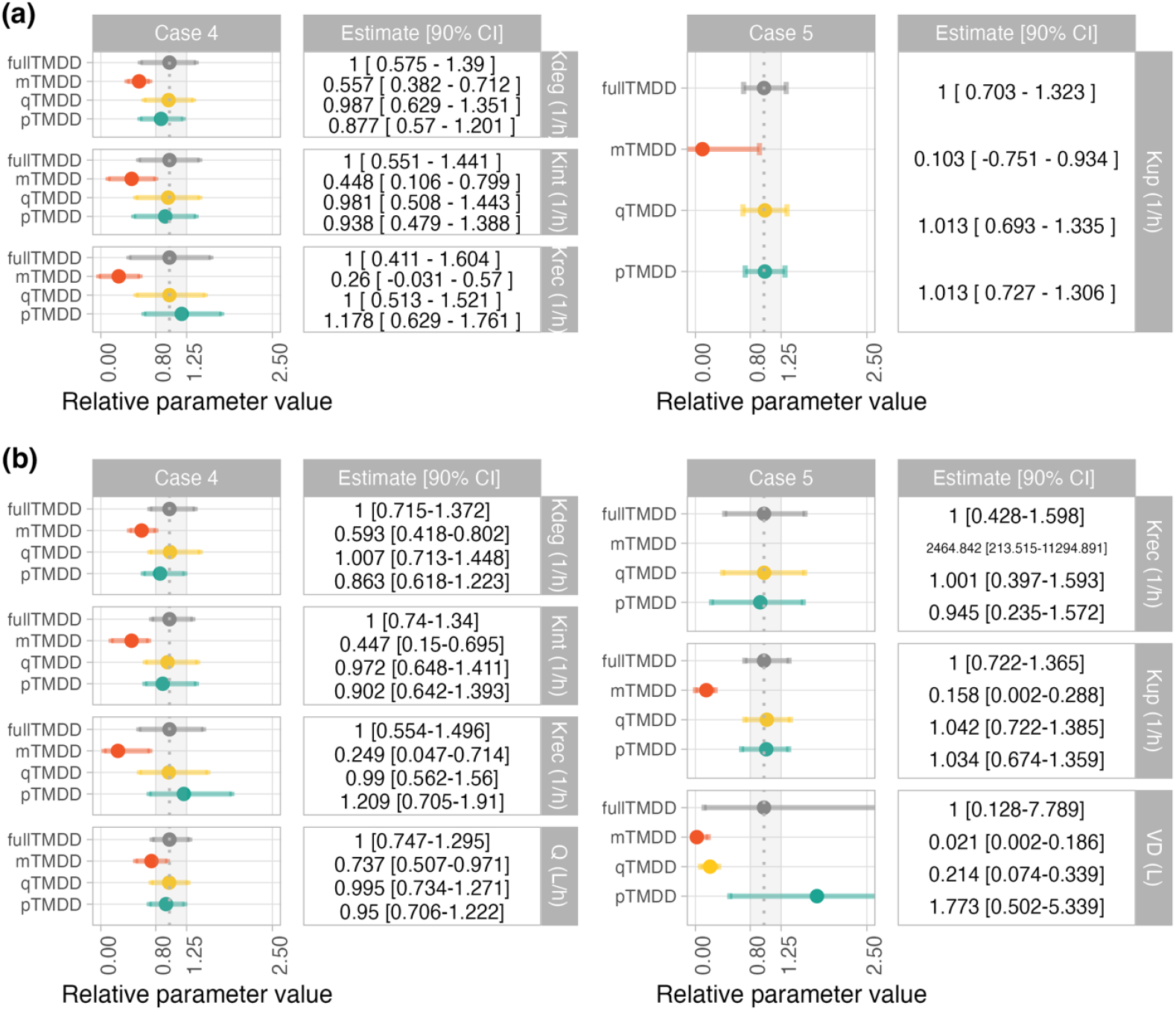
Impact of approximation methods on parameter estimate accuracy. (a) Parametric approach (b) non-parametric approach The closed dots represent the median of the predicted relative change from the reference model, and the 90% confidence interval (CI) associated with the medians was visualized using the error bars and includes the uncertainty in the predictions for the reference subject. The specific values of the medians and 90% CIs are shown in the statistics box on the right-hand side of the parameter; these values were calculated based on (a) 1000 sampled parameter vectors from the variance–covariance matrix obtained from NONMEM and (b) bootstrap results. The parameter values for a reference model (full TMDD) are shown by the dotted vertical line; the shaded area indicates the 80–125% margins relative to the reference model and are based on the standard bioequivalence limits. *k*_*deg*_: degradation rate of unbound drug, *k*_*int*_: internalization rate of the drug–target complex, *k*_*rec*_: FcRn-mediated recycling rate, *k*_*up*_: rate of cellular uptake in FcRn-mediated recycling space, *V*_*D*_: apparent volume of distribution of drug in FcRn-mediated recycling space, and *Q*: Inter-compartment clearance

Taken together, these results are consistent with those of the previous simulation study, indicating that pTMDD maintained a higher estimation accuracy than mTMDD, particularly under low-dose conditions.

The full TMDD model is the least approximated model and can be applied across all cases regardless of the validity conditions of approximation models. However, large number of parameters can lead to parameter identifiability issues, which may result in inflated %RSEs and reduced biological interpretability. This highlights the need for appropriately chosen approximation models. To quantify parameter uncertainty, we used Dixon’s Q test to count the number of parameters whose RSE was substantially larger or smaller than those of the other approximation models in each case.

Across both parametric and nonparametric assessments, mTMDD exhibited the largest number of parameters with substantially higher RSE. This discrepancy was observed primarily in cases where the V value was not much greater than 1. The largest difference was observed in Case 5, which was the only case in which *V*_2_fell below 1. In contrast, qTMDD and pTMDD consistently showed fewer parameters with elevated RSEs (Fig 2). Overall, parameter uncertainty was lower in the qTMDD and pTMDD model than in the full TMDD and mTMDD model.

**Fig 2.**
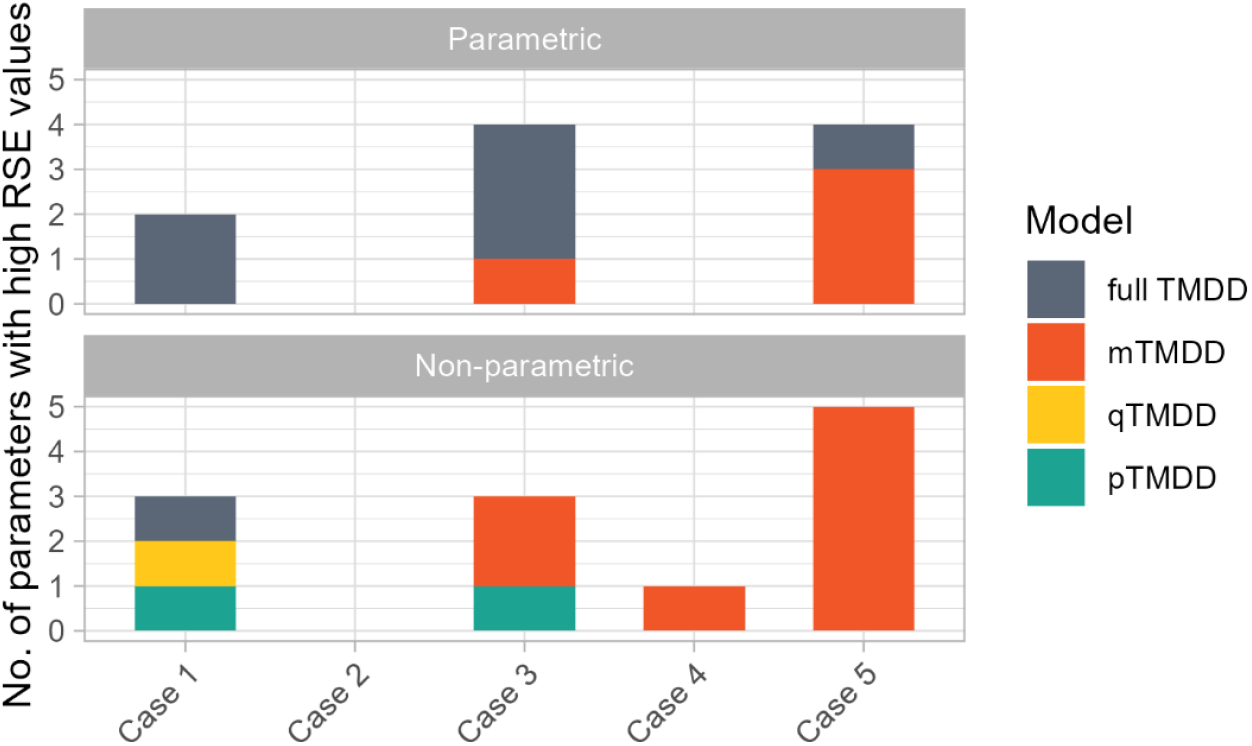
Number of parameters with high relative standard error (RSE) values. The values represent the number of parameters (out of all the estimated parameters) for which Dixon’s Q test indicated that the relative standard error was substantially larger than that of the other approximation models.

### pTMDD had shorter runtime than qTMDD

The computational efficiencies of the models were compared by measuring the elapsed time for parameter estimation and bootstrapping. For each case, the estimation time was defined as the average elapsed time for parameter estimation over five repeated runs, and the bootstrap time as the average elapsed time per replicate among the 1,000 bootstrap runs that achieved successful minimization and covariance steps.

For estimation time, the overall ordering of computational burden across models was full TMDD > qTMDD > mTMDD > pTMDD (Table 2a). For bootstrap time, the ordering was full TMDD > qTMDD > pTMDD > mTMDD (Table 2b). When considering the overall pattern across cases, pTMDD showed better computational efficiency than qTMDD for both estimation and bootstrap times.

**Table 2.** Elapsed time for all approximation models in each case.

| **(a) Estimation**<br>**time** | **full TMDD** | **mTMDD** | **qTMDD** | **pTMDD** |
| --- | --- | --- | --- | --- |
| **Case 1** | 371.6<br>[298.2–498] | 108.3<br>[77–139.9] | 63.2<br>[43.8–70] | 118.7<br>[84.3–154] |
| **Case 2** | 31.2 [30.8–34.5] | 5.9 [5.7–7.6] | 6.3 [6–6.8] | 5.6 [5.4–6.2] |
| **Case 3** | 51.5 [46–66] | 6.1 [5.7–6.5] | 9.7 [9.1–10.7] | 4.6 [4.2–5.1] |
| **Case 4** | 1470.8<br>[1301.4–1582.4] | 168.9<br>[146.4–189.9] | 340<br>[309–392.3] | 198.1<br>[175.3–223.8] |
| **Case 5** | 110.1<br>[95.6–129.5] | 78.4<br>[73.3–84.2] | 28.2<br>[22.3–35.4] | 17.8<br>[14.3–18.2] |
| **(b) Bootstrap**<br>**time** | **full TMDD** | **mTMDD** | **qTMDD** | **pTMDD** |
| **Case 1** | 28.1 [15.6–77.7] | 4.8 [2.1–93.9] | 5.6 [2.4–78.6] | 4.6 [1.9–26] |
| **Case 2** | 583.4<br>[209.6–3890.2] | 40.5<br>[15.4–199.8] | 43.5<br>[20.9–367.6] | 38.5<br>[16–428.2] |
| **Case 3** | 6127.6<br>[2347.8–20948.9] | 2179.7<br>[514.5–7551.5] | 2404.4<br>[642.5–6880.7] | 2311.7<br>[509.1–7279.5] |
| **Case 4** | 143.3<br>[63.4–450.2] | 14.7<br>[7.1–300.6] | 12.4<br>[6.3–34.8] | 9.7<br>[4.9–59.8] |
| **Case 5** | 166.6<br>[63.4–1353.5] | 116.9<br>[44.9–574.7] | 43.7<br>[21.1–180] | 53.7<br>[22.2–187.4] |

### Q2PCONV: R shiny application to convert qTMDD to pTMDD

To facilitate the practical use of pTMDD in routine PK analysis, an R shiny application that converts the NONMEM code to pTMDD was developed (https://github.com/hyese128/Q2PCONV). Although pTMDD is analytically simpler than qTMDD, direct implementation in NONMEM may be unfamiliar to many users. Q2PCONV addresses this problem by automatically translating qTMDD structures into pTMDD code while preserving the original model specification.

The Q2PCONV was validated using five clinical trial datasets and was found to work well for TMDD models with either a single receptor or two receptor pools (pharmacological receptor and FcRn). These results demonstrate that pTMDD is not only theoretically and empirically advantageous but also practically accessible through an automated model-conversion workflow.

## Discussion

Choosing an appropriate TMDD approximation method is critical because an inappropriate choice can, depending on the pharmacological setting, either greatly increase the computation time or compromise the parameter estimation accuracy. In this study, the advantages of using pTMDD were compared in three established TMDD models—mTMDD, qTMDD, and the full TMDD model—based on five clinical datasets from clinical trials. The evaluation was conducted from three perspectives: estimation, model diagnostics, and computational efficiency. These comparisons were interpreted in the context of previously proposed validity conditions for TMDD approximations. To relate these criteria to the clinical cases, they were quantified using two indices, V_1_ = (C_0_ + K_SS1_)/R_pharmacologic_ and V_2_ = (C_0_ + K_SS2_)/R_FcRn_, which separately captured TMDD through the pharmacological receptor and FcRn because most of the evaluated biologicals were IgG or Fc-fusion proteins that bound both targets.

For estimation results and model diagnostics, mTMDD showed the largest discrepancies in V_2_ in cases where these V_2_ values were clearly below 1, whereas qTMDD and pTMDD remained closer to the full TMDD model. This was consistent with our hypothesis that pTMDD would be especially effective under conditions where the drug concentration was low relative to the receptor capacity. Although TMDD is often considered relevant at high drug concentrations, it must be accurately described at low doses. At low doses, the PK profile may appear linear, and TMDD models are frequently not considered; however, elimination in this range can still be governed by drug–target interactions and only appears linear. Misinterpreting such patterns as simple linear elimination can lead to an incomplete understanding of drug disposition, for example, failing to explain a rapid initial decline in plasma concentrations due to extensive target binding or an apparent drop in systemic levels caused by rapid distribution into tissues with high target expression. Therefore, approximation models for TMDD should remain valid across a wide range of dose levels and receptor capacities, including conditions in which C_0_ + K_SS_ is not significantly greater than R_tot_.

The estimation time was shortest for pTMDD in most cases, with particularly large reductions in complex scenarios. For instance, in Case 4, the computation time decreased from 1,447 s with the full TMDD model to 345 s with qTMDD and 200 s with pTMDD. The improved computational efficiency of pTMDD relative to qTMDD is expected to facilitate population analyses of large clinical datasets, thereby saving time and computational resources. It also enables extensive scenario exploration, including a wide range of doses, dosing intervals, and patient subgroups, ultimately supporting more efficient and informed decision making throughout the model-informed drug development process.

One limitation of the study is that, except for special situations such as microdosing, drug development studies typically use doses that greatly exceed receptor levels, resulting in *V* > 1 in most cases, which makes it difficult to clearly demonstrate the differences in parameter accuracy between approximation models. Additional analyses in settings where *V* ≤ 1 would therefore be valuable to further substantiate the conclusions.

## Materials and Methods

### Selection of population pharmacokinetic models

Five population PK models based on clinical trial data (NCT02175056, NCT03644056, NCT04414163, NCT05276310, NCT02860715) were selected[8-11], and approximation models were selected for analysis. All models considered inter-individual and residual variability and were selected based on numerical and visual criteria.

For each dataset, four model versions were implemented: the full TMDD model, mTMDD, qTMDD and pTMDD. Apart from the TMDD approximation part, all other structural elements of the models were kept identical to the original model code. To enable model comparison, the total receptor concentration (R_tot_), neonatal Fc receptor concentration (FcRn), equilibrium constant for the drug-FcRn interaction (K_SS1_), and equilibrium constant for the drug-receptor interaction (K_SS2_) were fixed. All the other parameters were estimated.

### Comparison of parameter estimation results

Parameter estimation for all model variants was performed using the first-order conditional estimation with interaction (FOCE-I) method in NONMEM 7.5, with PsN 5.3.1. For each approximation model, the final estimates from the corresponding full TMDD model were used as initial values to minimize the effect of initialization.

#### Comparison of parameter estimate accuracy and model diagnostics

To compare estimation results, the results of the full TMDD model were set as reference values and the closeness of parameter estimates from each approximation model to the reference was evaluated. Both parametric and non-parametric assessments were conducted, following the methodology of Jonsson and Nyberg [13]. The parametric approach was based on the variance–covariance matrix obtained from NONMEM. cov file, whereas the non-parametric approach was based on bootstrap results.

Parameter estimate accuracy was assessed by comparing the quantiles of the 1,000 sampled parameter vectors from each approximation model with the corresponding median value from the full TMDD model. Medians and 5^th^-95^th^ percentiles were compared with the median estimates from the full TMDD model. Visualization and comparison of fixed-effect parameters were performed using the PMXForest R package. An equivalence range of 80%-125% of the full TMDD median was used as a reference interval, consistent with standard bioequivalence criteria [13].

Model performance was evaluated using standard diagnostic criteria in population PK modeling. Model fit was assessed using the OFV and GOF plots. The model reproducibility was examined using VPC plots. The diagnostic results of each approximation model were compared with those of the full TMDD model.

#### Comparison of parameter uncertainty

The parameter uncertainty was also evaluated using both parametric and non-parametric approaches by comparing the RSE values across the models. The RSE was calculated as:

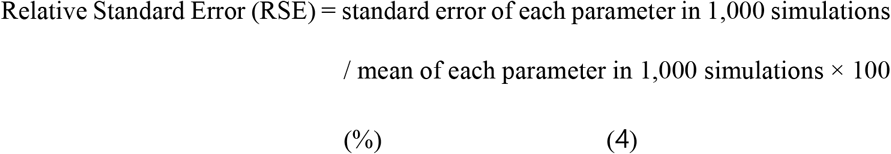

In this assessment, Dixon’s Q test was applied to identify whether the RSE value of a specific parameter from a given approximation model was substantially higher than those of the others. Models with substantially elevated RSEs as determined using this test were considered to have poor estimation precision.

### Comparison of computational efficiency

To evaluate computational efficiency, both the estimation and bootstrap runtimes were assessed. The estimation time was quantified based on the elapsed estimation time reported in the .lst files from five independent runs of the final model. For bootstrap analysis, 1,000 replicates were generated using the final model, and only runs with successful minimization were included in the comparison of the bootstrap times.

The system used for the computation was an IdeaPad 5 Pro 16ACH6 equipped with 8GB RAM and an AMD Ryzen 7 5800H (3.2 GHz) processor. To ensure a fair comparison between the approximation models, all the models were evaluated under the same settings with consistent $SUBROUTINE options (e.g., ADVAN, TOL) and $EST options (e.g., NSIG, SIGL).

### Development of Q2PCONV

Q2PCONV, an R Shiny application, was developed to facilitate a fast and smooth transition from the qTMDD to the pTMDD model. The Shiny package in R is a tool for building interactive web applications that enables easy data visualization and analysis in a web environment. It supports real-time data updates and automated computations, thereby enhancing workflow efficiency. Q2PCONV is fully compatible with existing NONMEM model codes and features a user-friendly interface that allows users to input and modify the code with maximum flexibility. This allows researchers to translate a qTMDD model into a pTMDD model with greater ease and efficiency.

## Data Availability

All relevant data are within the manuscript and its Supporting Information files.

## Code Availability

The NONMEM code of the population pharmacokinetic model with each approximation method is available in S1 Text.

## Acknowledgements

This study was supported by Chungnam National University, Institute of Information & Communications Technology Planning Evaluation (IITP) grant funded by the Korea government (MSIT) (No.RS-2022-00155857, Artificial Intelligence Convergence Innovation Human Resources Development (Chungnam National University)), National Research Foundation of Korea (NRF) grant funded by the Korea government (MSIT; No. RS-2023-00278597, RS-2022-NR069643, RS-2022-NR070856, Senior Health Convergence Research Center based on Life Cycle (Chungnam National University)), Korea Environmental Industry & Technology Institute (KEITI) through Core Technology Development Project for Environmental Diseases Prevention and Management (RS-2021-KE001333), funded by the Korea Ministry of Environment (MOE), a grant of the Korea Machine Learning Ledger Orchestration for Drug Discovery Project(K-MELLODDY), funded by the Ministry of Health & Welfare and Ministry of Science and ICT, Republic of Korea (grant number : RS-2024-00460694), the Korea Institute of Toxicology (KIT) Research Program (no. 2710008763, KK-2401-01), Korea Health Technology R&D Project through the Korea Health Industry Development Institute (KHIDI) grant funded by the Ministry of Health & Welfare, Republic of Korea (grant number : RS-2024-00336984, RS-2025-02306055), supported by the Ministry of Trade, Industry, and Energy (MOTIE), Korea, under the “Infrastructure program for industrial innovation” supervised by the Korea Institute for Advancement of Technology (KIAT)(RS-2024-00434342), Basic Science Research Program through the National Research Foundation of Korea (NRF) funded by the Ministry of Education (RS-2025-25397599), BK21 FOUR Program by Chungnam National University Research Grant, 2024.

## Author Contributions

Conceptualization: Hwi-yeol Yun, Jae Kyoung Kim, Jong Hyuk Byun

Data curation: Hyeseon Jeon, Woojin Jung

Formal analysis: Hyeseon Jeon

Funding acquisition: Hwi-yeol Yun, Soyoung Lee, Jae Kyoung Kim, Jung-woo CHAE, Jong Hyuk Byun

Investigation: All authors

Methodology: Hwi-yeol Yun, Soyoung Lee, Jung-woo CHAE

Project administration: Hwi-yeol Yun, Soyoung Lee, Jung-woo CHAE

Supervision: Jae Kyoung Kim, Jong Hyuk Byun

Validation: All authors

Visualization: Hyeseon Jeon, Woojin Jung

Writing – original draft: Hyeseon Jeon

Writing – review & editing: All authors

## Supporting information captions

**S1 Table. Objective Function Value (OFV) and Akaike information criteria (AIC) values of final approximation models for each case**

**S2 Table. NONMEM dataset for all cases**

**S1 Fig. Visual Predictive Check results for each approximation model in each case**

The blue shading indicates the model-predicted confidence intervals of the 5th and 95th percentiles, and the red shading indicates the model-predicted confidence intervals of the 50th percentile. For all pharmacokinetic models, the model-predicted confidence intervals encompassed the 5th, 95th, and median observations.

**S2 Fig. Basic goodness-of-fit plots for each approximation model in each case**

(A) Population prediction vs. observation; (B) individual prediction vs. observation; (C) individual prediction vs. CWRES; and (D) TIME vs. CWRES. Plots (A) and (B) showed trends close to the line of identity (*y* = *x*), whereas CWRES values in plots (C) and (D) were mostly from −2 to 2. No clear differences were observed between the approximate models across all the diagnostic plots. TMDD, ; mTMDD, ; qTMDD, ; pTMDD,

**S1 Text. The NONMEM code of the population pharmacokinetic model with each approximation method implemented**.

